# Finding Hidden Treasures: A Child-Friendly Neural Test of Task-Following in Individuals Using Functional Transcranial Doppler Ultrasound

**DOI:** 10.1101/815910

**Authors:** Selene Petit, Nicholas A. Badcock, Alexandra Woolgar

## Abstract

Despite growing interest in the mental life of individuals who cannot communicate verbally, objective and non-invasive tests of covert cognition are still sparse. In this study, we assessed the ability of neurotypical children to understand and follow task instructions by measuring neural responses through functional transcranial Doppler ultrasound (fTCD). We recorded blood flow velocity for the two brain hemispheres of twenty children (aged 9 to 12) while they performed either a language task or a visuospatial memory task, on identical visual stimuli. We extracted measures of neural lateralisation for the two tasks separately to investigate lateralisation, and we compared the left-minus-right pattern of activation across tasks to assess task-following. At the group level, we found that neural responses were left-lateralised when children performed the language task, and not when they performed the visuospatial task. However, with statistically robust analyses and controlled paradigms, significant lateralisation in individual children was less frequent than expected from the literature. Nonetheless, the pattern of hemispheric activation for the two tasks allowed us to confirm task-following in the group of participants, as well as in over half of the individuals. This provides a promising avenue for a covert and inexpensive test of children’s ability to covertly follow task instructions and perform different mental tasks on identical stimuli.

## 1 Introduction

Modern neuroscience is taking a growing interest in the mental life of individuals who may not be able to overtly display the extent of their cognitive abilities. In the case of vegetative patients, or minimally verbal autistic individuals ^1^, for example, recent evidence has suggested intact consciousness and language comprehension, despite an absence of communicative behaviour (e.g., Cruse et al., 2011; Owen et al., 2006 for minimally-conscious patients, and Cantiani et al., 2016; DiStefano et al., 2019; Kedar, 2012 for minimally-verbal autistic individuals). In minimally verbal autism, in particular, it appears that cognitive abilities may be under-estimated by standard assessments, due to inability to comply with task-demands, lack of motivation, or demanding social constraints associated with the testing situations (Kasari et al., 2013). For this reason, it is crucial to develop a reliable test of covert cognitive abilities that does not rely on behavioural responses. In this study, we aimed to develop such a method, using a portable and easy-to-setup neuroimaging technology, and to validate the method with typically-developing children. We used the logic that if we could index task-following directly from neural signals, this could be used to assess language comprehension and other cognitive abilities.

We took our inspiration from previous research that used functional neuroimaging to study cognitive abilities in non-communicative patients. In a seminal study, Owen et al. (2006) instructed a patient in vegetative state to perform one of two mental imagery tasks (imagining playing tennis or imagining walking around her house). The patient’s brain responses measured with magnetic resonance imaging (fMRI) were significantly different between the two conditions, suggesting that the patient was able to understand the instructions and wilfully follow the task commands. These results have been replicated and expanded in several studies requiring patients to follow different instructions, such as imagining moving their right versus left hand (Bekinschtein et al., 2011), counting versus listening to words (Monti et al., 2009), or naming pictures (Rodriguez Moreno, Schiff, Giacino, Kalmar, & Hirsch, 2010). The logic of these studies indicates that task-following may be a useful index into the mental life of individuals who do not otherwise communicate.

The high cost of MRI, the requirements to lie still in the scanner, and the noise associated with the scanning procedure make this method inaccessible to some populations such as young children and some autistic individuals. However, recently, research teams have begun to use functional transcranial Doppler ultrasonography (fTCD) as a non-invasive and relatively inexpensive alternative to fMRI (Lohmann et al., 2006). Being relatively insensitive to movements, fTCD allows for testing a wider range of populations, including those with difficulties staying still such as children (Lohmann et al., 2006) and infants (Kohler et al., 2015), and it has previously been used in populations where standard language assessments may not be suitable such as deaf children (Payne et al., 2019). It is also portable, allowing it to be used outside of the laboratory, and in larger populations. FTCD uses two probes placed on participants’ left and right temples to measure the blood flow velocity through the left and right middle cerebral arteries. It is inferred that faster blood flow to one hemisphere results from higher neural activity in that region. Thus, fTCD allows for an indirect measure of brain activation in the two hemispheres, and can be used to examine the lateralisation of neural responses associated with different cognitive processes.

Our aim was to derive an implicit measure of language comprehension that could be used in non-speaking populations such as minimally-verbal autistic children. We combined the logic of task-following paradigms, in which evidence for wilful modulation of neural activity must reflect comprehension of verbal instructions, with the accessible technology of fTCD. In particular, we aimed to use fTCD to provide a measure of differential brain activation in response to different tasks. We employed two tasks that primarily elicit activity in the left and the right hemispheres respectively: a word generation task and a visuospatial memory task (Bishop et al., 2009; Groen et al., 2012; Rosch et al., 2012). During both tasks, participants were presented with a spatial array in which a single letter was presented in several locations. In the word generation task, participants were asked to silently generate as many words as possible starting with this letter. During the visuospatial memory task, participants were asked to study the location of letters and remember their location after the letters disappeared.

We chose to compare lateralisation between two tasks, rather than using a single task, based on the observation that lateralisation of language and visual-spatial processing varies across individuals, but still tends to be complimentary (i.e., different for the two tasks) across individuals. For example, current estimates are that around 7.5% to 25% of the population have right hemisphere language and around 10% to 15% have bilateral representation of language functions, with the remaining 60% to 80% having the typical left representation of language (Knecht et al., 2000; Lust et al., 2011; Whitehouse & Bishop, 2009). Similarly, visuospatial functions are not supported by the right hemisphere in every individual (Badcock, Nye, et al., 2012; Rosch et al., 2012). In their respective studies, Whitehouse and Bishop (2009) found that 25% of adults had either a bilateral or a left-hemisphere dominance for visuospatial memory, while Groen et al. (2012) found this pattern in 29% of children. In addition, individual factors such as handedness may influence lateralisation in individuals (Groen et al., 2012). Thus, for our purpose of measuring task-following on an individual-subject basis, it may be difficult to interpret the result from a single task. However, the major theories of lateralisation do nonetheless converge on the idea that language and visuo-spatial functions are supported by separate hemispheres, to optimize cognitive performance (Cai et al., 2013; Heilman et al., 2000; Whitehouse & Bishop, 2009). For example, using fMRI, Cai et al. (2013) found complementary lateralisation of language and spatial attention for all but one of 29 participants. The origin of this distribution might be causal (Cai et al., 2013; Heilman et al., 2000), i.e., one function is lateralised to one hemisphere because the other is lateralised to the opposite hemisphere, or result from independent biases (Badzakova-Trajkov et al., 2016; Knecht et al., 2000; Whitehouse & Bishop, 2009; Zago et al., 2016). These two major theories differ in their explanation of the origin of complementarity, but converge on the expectation that hemispheric complementarity exists in the majority of individuals. Thus, our study was based on the logic that if the two functions rely on different hemispheres in most individuals, it should be possible to observe differential task-related activity in individual children. In addition, the fact that the language task is inherently a productive task, while the visuo-spatial memory task is a receptive task is advantageous for our purpose of measuring differential lateralisation between tasks as the neural patterns associated with these tasks will likely unfold over different timeframes (Bishop et al., 2009; Groen et al., 2012). Moreover, directly comparing the two tasks allowed us to be sensitive both to differences in the lateralisation and to differences in the time course with which the lateralisation occurs. For example, if the two tasks are both lateralised to the left hemisphere for a particular individual, but the change in velocity happens more quickly for word generation, this would manifest in a measurable difference in the lateralisation between tasks at early time points, even if both tasks are ultimately lateralized to the same hemisphere.

In developing our approach, we addressed two limitations that would otherwise prevent the clinical application of this method as a test of task-following. First, an issue in extant fTCD research is the heterogeneity in paradigms used to measure different cognitive processes. For instance, most researchers estimate language lateralisation using word generation paradigms that involve generating language after viewing a letter on a screen or a short animation (Badcock, Nye, et al., 2012; Rosch et al., 2012; Woodhead et al., 2018). On the other hand, most researchers estimate visuospatial lateralisation using paradigms with complex visual displays, such as finding rabbits hiding in different holes, or lines masked by complex visual dynamic masks (Groen et al., 2011; Rosch et al., 2012). As such, the difference in lateralisation between tasks may correspond to changes in the visual and auditory stimuli instead of differences in language and visuospatial processes. Thus, a secondary aim of the research was to report lateralisation effects for language and visuospatial tasks using identical visual stimuli.

The second limitation concerned a statistical flaw in that the way that lateralisation indices (LIs) are commonly analysed, which systematically over-estimates laterality. Typically, LIs are calculated by finding the peak in the left-right blood flow velocity difference, then averaging the velocity values over a time-window (usually 2 s) centred on that peak (e.g., Badcock, Nye, et al., 2012; Deppe, Knecht, Lohmann, & Ringelstein, 2004; Groen et al., 2011; Kohler et al., 2015; Woodhead et al., 2018). This quantifies the *maximum* difference of the waveform, which can then be compared between tasks groups. However, because of the way it is derived (taking a maximum from a continuous waveform), it is statistically biased to compare this difference to zero, for example, to infer that the group or individuals have “significant” lateralisation. We show through simulation that this method, which is common practice in fTCD research, will tend to push individual LIs away from zero, inflating type I (false positive) error. This flaw, also known as ‘double dipping’, is well known in fMRI and electroencephalography research (Kilner, 2013; Kriegeskorte et al., 2009) and applies equally to fTCD data. We show through simulation that it can be avoided by omitting the peak selection (i.e., averaging over the entire *a priori* period of interest), yielding LIs that can legitimately be compared to zero to infer significance of lateralisation. We employ this statistically-robust approach for our analysis.

We hypothesised that if children consistently performed the language and visuo-spatial memory tasks as instructed, we would observe distinct hemispheric patterns of activation between two mental tasks, both at the group level and in individual children. Using our controlled stimuli and statistically unbiased analyses, we found robust evidence that the two tasks relied on different brain processes, as expected, which was indicated by a difference in the pattern of hemispheric activation in the group. However, at the individual level our sensitivity was medium, with statistical evidence of task-following detected in only 55% of children. This could reflect a tendency for some children to ignore task instructions, or reflect a relative insensitivity of the approach. The work provides a possible avenue for a covert and inexpensive assessment of task-following in children, but individual sensitivity would need to be improved for clinical application.

## 2 Methods

All presentation scripts, analysis scripts, and raw data are available at https://osf.io/xygjv/.

### 2.1 Participants

Twenty-two children were recruited using the Neuronauts database of the Australian Research Council Centre of Excellence in Cognition and its Disorders. All participants were native English speakers, and they received $20 for their participation. The data from two participants were excluded due to failing to record data (one participant) and computer crashing (one participant). The final set of data thus came from 20 participants (age range: 9 to 12 years old, *M*=*10:7*, *SD*=*1:1,* 10 male and 10 female). Seventeen of the participants were right-handed, and three were left-handed, based upon parent reports. This study was approved by the Macquarie University Human Research Ethics Committee (Reference number: 5201500074). Participants’ parents or guardians provided written consent and the children provided verbal consent.

### 2.2 Apparatus

We acquired blood flow velocity data using a Doppler ultrasonography device (Delica EMS-9UA, SMT medical technology GmbH&Co Wuerzburg, Germany), with probes held in place bilaterally over the left and right temporal windows via a headset. We adjusted the probes until we obtained a good signal of the blood flow through the left and right middle cerebral arteries. The experimental paradigm was presented using Psychtoolbox version 3 (Brainard, 1997; Kleiner et al., 2007) on Matlab, on a 27-inch monitor screen located at 80cm from the participants. Responses to the trials were given via a button box (Cedrus RB-830).

### 2.3 Paradigm

In order to engage children with the task, we presented the paradigm as a game in which the children collected treasure. A male and a female “pirate” gave auditory instructions and feedback on each trial. The pirates’ voices were recorded by actors who were native Australian English speakers. Male and female voices were included for diversity and were not related to the two tasks (both voices instructed both tasks with equal probability). Each participant completed 40, one-minute trials, switching tasks every 10 trials. They completed 20 trials of the word generation task (task 1), and 20 trials of the visuospatial memory task (task 2). The order of the task was counterbalanced across participants.

Each trial started with a baseline period of 10 s, during which the participants fixated on a black cross in the centre of a white screen. Then either the male (first half of the experiment), or the female (second half of the experiment) pirate was presented on screen, and greeted the participant. The pirate asked the participant to get ready, and gave the instructions for the task. The instructions were “Think of words that begin with this letter” for task 1, and “Remember my treasure map” for task 2. Then a treasure map appeared, with 8 repetitions of a letter randomly distributed on the screen (see Figure 1). The characters were displayed in black, presented at a visual angle of approximately 1°. The treasure map remained visible for 5 s during which children silently generated words (task 1) or studied the position of the letters (task 2). A white screen was then displayed for 10 s, during which the children continued to generate words (task 1) or remembered the position of the letters (task 2). Finally the map reappeared with the letters either exactly at the same location as the first map (in half of the trials), or with one letter displaced from its original location. The pirate would then ask “Did you think of lots of words?” (task 1) or “Is this the same treasure map?” (task 2), and the children would answer “yes” or “no” by pressing either a right or left button on a button box in front of them. The position of the buttons was counterbalanced across participants. Contrary to most fTCD studies, we chose not to ask participants to overtly name the words that they generated, as we had designed the paradigm to eventually be used for non-speaking individuals. A 5 s animation was then presented, showing the treasure that the pirate collected during the trial, with the pirate giving encouraging auditory feedback (e.g., “You’re winning!”, “Blow me down, that was brilliant!”, “Pieces of eight, you’re doing great!”). Then the pirate’s voice would indicate that the child should take a break by saying e.g., “Time for a rest” and a short animation showed a relaxing situation in which the pirate was yawning or sailing away for the night. This was included to encourage participants to stop performing the tasks, with the intention of encouraging task-related activation to return to baseline. Finally, a blank white screen was presented for 10 s of normalisation, then the next trial began. Each trial featured a different letter, with all the letters of the Roman alphabet being presented once in each task, except for the letters K, Q, W, X, Y, and Z, which were not used as words starting with these letters are rare. The order of the letters was randomized for each participant. Each letter was seen once in a single task before being repeated in the other task. The order of the letters was reversed in the second task, for each participant. Thus, the paradigm was designed so that the two tasks consisted of identical visual stimuli and identical structure, and differed only in auditory instructions. Any difference in the hemispheric activation must therefore be attributed either to the subtly different auditory stimulation, or to the difference in the mental task.

**Figure 1:**
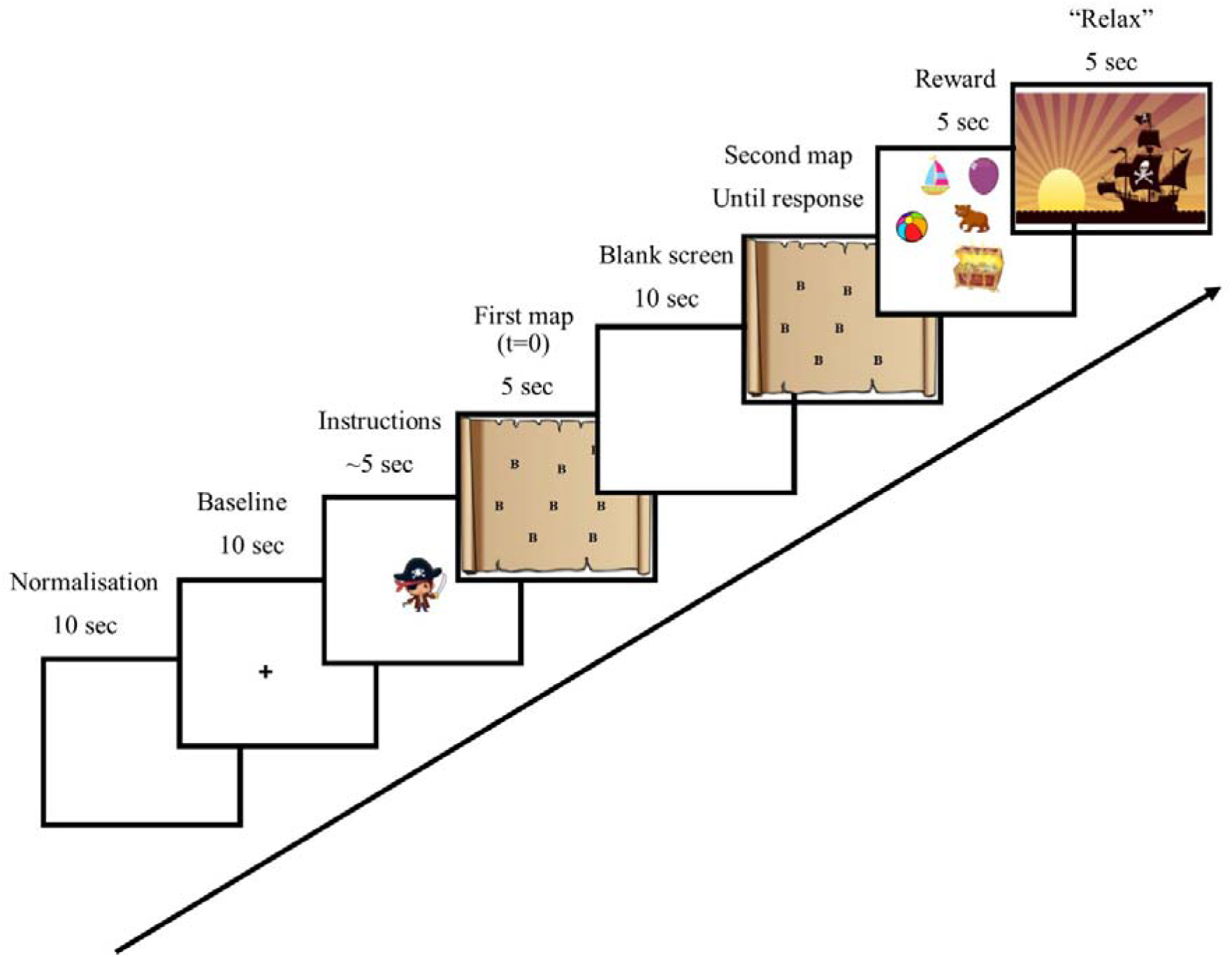
Trial structure. After normalisation and baseline, a pirate was presented and introduced the task. Then a treasure map with letters appeared and children started generating words (language task) or remembering the location of the letters (visuospatial memory task). The map then disappeared, and reappeared with all letters at the same location, or with one letter that changed location. A reward screen then appeared, followed by an animation instructing children to relax.

### 2.4 Data pre-processing

We pre-processed the fTCD data using DOPOSCCI (Badcock et al., 2018; Badcock, Holt, et al., 2012) with MATLAB version R2017B (Mathworks Inc., Sherborn, MA, USA). We first down sampled the raw data to 25Hz, then we removed the heart cycle by determining local peaks and using linear heart cycle correction based on previous work (Badcock, Nye, et al., 2012). To correct for overall differences in the strength of the signal from the right and left probe (e.g., due to a difference in the alignment of the probes), we normalised the signals to a mean of 100% on a trial-by-trial basis. We then created epochs, −15 to 40 s, relative to the onset of the first visual map display (see Figure 1). At this stage, we rejected epochs with extreme values (beyond ± 50% of the mean signal), corresponding to poor insonation or excessive head movement. Finally, we performed a baseline correction for each epoch by removing the averaged value of the signal from −15 to −10 s before stimulus onset.

### 2.5 Lateralisation Index

For comparison with previous literature, and to test for differences in lateralisation between two tasks using similar stimuli, we calculated lateralisation indices (LIs). Typical fTCD lateralisation research calculates LIs by finding the peak in the mean velocity difference between the two hemispheres within a POI, extracting data from a 2s time-window around this peak, averaging over the window, and comparing this value to zero. This approach provides a metric that can be compared between groups or conditions, but, because of the way it is derived, it should not be compared to chance at the individual or group level. This is because the time-window is selected to include the peak in the data, so even if there is no effect (random noise) the average value will tend to be different from zero. Given that there are many observations to choose from (the entire time course) selecting the time-window in this way will increase false positive rate, an effect identified in other disciplines as “double dipping” (Abbott, 2009; Kilner, 2013; Kriegeskorte et al., 2009). This effect is mitigated by averaging over a large time window (2s), but, we argue, will not be completely removed. We first simulated the problem and a solution, and then applied the statistically robust approach to our empirical data.

#### 2.5.1 Simulation of statistically-robust method for determining LI

To confirm our intuition that the typical approach (“peak selection” method) to LI derivation in the literature will increase false positives, we ran the analysis on simulated random data (no true signal). We compared the results to those from a statistically robust alternative approach in which there is no data selection: signals are averaged over an entire, pre-defined, POI (“no selection” method). The simulation was as follows: for each hypothetical “individual”, we generated twenty noisy time-series (“trials”) of 200 time points (approximately the size of our language POI) by selecting a random number (“lateralisation”) between −20 and 20 at each time point (Figure 2A, grey lines). We then averaged each of these trials to create an “individual-subject” level mean (Figure 2A, solid black line). Proceeding as in the fTCD literature (“peak selection” method), we then found the peak in individual’s trial-averaged data and defined our POI as a 2 second (50 time points) time window centred on the peak. For the statistically robust “no selection” approach, we chose the entire time window as our POI (randomly sampling a subset of 50 values to make it comparable to the peak selection approach). For each method, we extracted the mean value across time in the POI for each of the 20 trials and considered their distribution. As we anticipated, the distribution of lateralisation over trials was shifted away zero for the peak method. This is illustrated for two “individuals” in Figure 2, one for whom the peak in the random noise was positive (Figure 2C) and one for whom it was negative (Figure 2G). As there is no true signal in the simulated data, these distributions should be centred on zero, as they are in the “no selection” method (Figure 2D and 2H).

**Figure 2:**
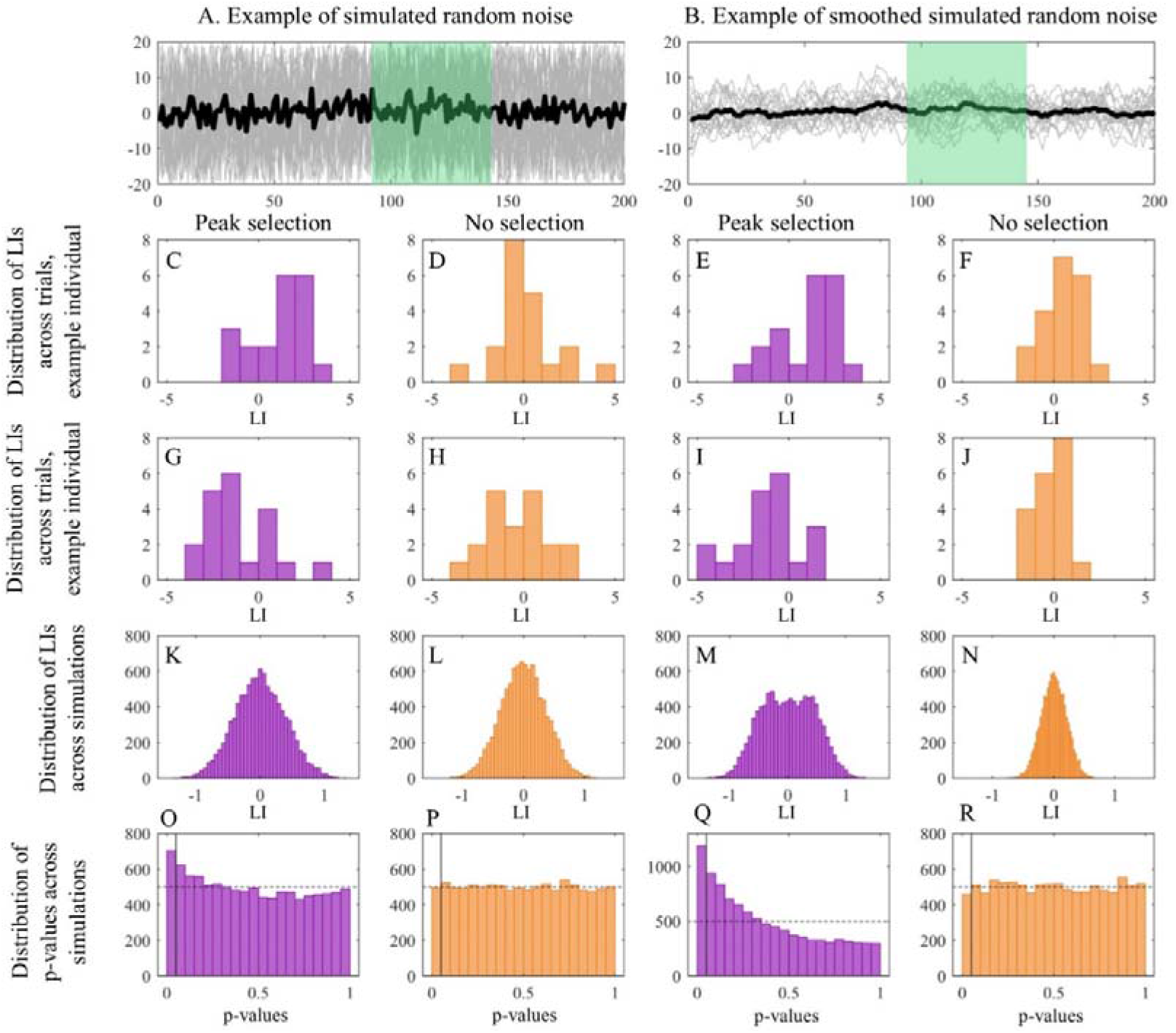
Simulation of LI significance using the “peak selection” (purple) and “no selection” (orange) methods. We simulated 20 “trials” of 200 time-points by randomly selecting a number between −20 and 20 at each time point (A, grey lines). We then averaged these to give an “individual subject” trial-averaged time-series response (A, black line). We then either found the peak in this averaged time-series and selected a 50 time points time-window around that peak (A, green area) to illustrate the peak selection approach, or randomly selected 50 time points to illustrate the no selection method. The resulting distribution of lateralisation for the 20 trials for an example individual is shown in the second row (C and D). Using the peak method, this distribution was shifted positively from zero (C), while using the average method it was centred on zero (D). An example of another simulated individual is shown in the third row. This individual shows a negative shift of lateralisation for the peak analysis (G), but data are again centered on zero for the average analysis (H). We generated 10,000 simulations, and then computed the LI of each simulation by averaging the data during the selected time-window, and plotted the distribution of these 10,000 LIs for method (K and L). Finally we compared each individual subject’s LI to zero in a two-tailed t-test and plotted the distribution of p-values across simulations (O and P). The peak analysis drastically increased the false-positive error rate (at α = .05, 500 simulations should fall below .05, O), while the average analysis did not (P). We further illustrate that this problem is likely to be exacerbated in actual fTCD data, by repeating the same procedure after temporally smoothing the data (right half of the Figure, B, E, F, I, J). In addition to increasing the false-positive error-rate, the bimodality in the distribution of LIs across the group is more visible (M).

In fTCD research, the mean across these trials would be taken as the LI for an individual, and the distribution across trials could be compared to zero to test it for significance. To illustrate why this is problematic, we ran the above simulation 10,000 times, and calculated an LI for each of them. As can be seen in Figure 2, the distribution of LIs across the population tends to be broader (more extreme lateralisation in either direction) for the peak selection method (Figure 2K) compared to the no selection method (Figure 2L). Most critically, we computed a two-tailed t-test of the LI against zero (no information in the signal) for each simulation, equivalent to testing the significance of the LI in an individual. The distribution of p-values is shown in the final row of Figure 2. If the test were statistically robust, we would expect 500 of the 10,000 simulations (5%) to have a p-value of 0.05 or less. Instead, for the peak method, the distribution is skewed towards small p-values (Figure 2O). This demonstrates a statistical bias to reject the null hypothesis (find a p-value of <0.05) more than 5% of the time. The effect was again not present using the “no selection” approach (Figure 2P). Finally, to quantify the problem, we calculated the percentage of simulations for which the LI was significantly different from zero, at alpha = 5%, *i.e.,* the false positive error rate. The peak method yielded an inflated false positive error rate of 7.1%, while the average method yielded a false positive error rate of 5%.

Note that this simulation most likely underestimates the problem in real fTCD data, because it will increase with smoothness in the timeseries data. To illustrate this, we performed the same procedure on the same simulations after temporally-smoothing the individual-trial time-series (Figure 2, right half). As can be seen, the problem becomes much more extreme when data are smooth, yielding now false-positive detection rate of 11.9% for the peak selection method (Figure 2Q), while the false positive rate for the no selection method stays close to the chosen alpha level at 4.6% (Figure 2R). In this simulation the effect on the distribution of LIs across the group is even more apparent: since each individual’s peak is shifted away from zero, and this shift can be in either direction, the result is a bimodal distribution at the group level (Figure 2M). Similar results have been found in real fTCD data by Woodhead et al. (2018), who found a bimodal distribution of lateralisation when analysing their data using the peak selection method, but a normal distribution when using the no selection method.

Note also that although we conceptualised the first level of our simulation as “trials” and the second level as “individuals” and tested the effect at the “individual-subject” level, the exact same result obtains if the first level were to be considered “individuals” and the peak selection and analyses were to be carried out at the “group” level. The statistical bias introduced for by the peak selection would then increase the false positive rate in the group level analysis.

#### 2.5.2 Treatment of empirical data

For our empirical analysis, we calculated the LI using the statistically robust “no selection method”. For each task separately, we first averaged the signal from all the accepted epochs, for the right and the left probes. We then calculated the difference between the left and right signals over time. We defined a language POI as 4 to 14 s after the first map onset, in accordance with previous research that found the highest left-hemisphere activation during this POI for the word generation task (Bishop et al., 2009; Groen et al., 2011, 2012). We defined a visuospatial POI as 20 to 35 s after the first map onset based on previous findings of highest right-lateralisation during this time-window (Groen et al., 2011, 2012; Rosch et al., 2012). For each task, we assessed the left-minus-right signal difference within the corresponding POI using a grand average within the POI and performing a one-sampled t-test between this difference and zero. This was done at the group level (across participants) and at the individual level (across trials within participants). We additionally calculated Cohen’s d (effect size).

At the request of reviewers, we also include a follow-up analysis using the statistically-biased “peak selection” method, selecting a 2s time window around the trial-averaged peak within each POI for each individual, to illustrate how this procedure would change the empirical result.

### 2.6 Split-half reliability

In addition to reporting the LIs for the group and individuals, we estimated the reliability of the LIs by calculating the split-half reliability for each task. This was done using Pearson’s correlation between the LI of each participant for the odd and the even trials, and was carried out for the statistically robust version of LI derivation only. We found good reliability for the word generation task (r = .58, p = .0068), and for the visuospatial memory task (r = .75, p < .001).

### 2.7 Hemispheric differences analyses

Finally, to address our main question of whether we could measure differential patterns of hemispheric activation between the two tasks, we compared the left-minus-right difference in blood flow velocity between tasks. We performed a two-tailed paired-sample t-test for the average blood flow velocity within the language task period of interest (POI). This POI was chosen as it has showed the strongest lateralisation for language, and no lateralisation for visuospatial processing, in previous research (Badcock, Nye, et al., 2012; Groen et al., 2012; Whitehouse & Bishop, 2009), so we expected the lateralisation for the two tasks to be maximally different during this period. By only analysing the left-right difference once (in just the language POI), we avoid the need to correct for multiple comparisons thus maximising our statistical power. We performed this analysis at the group level, with a paired t-test across participants, and at the individual level, with a paired t-test across trials (pairing the letters in each condition) within participants. At the group level, with 20 participants, we had .56 power to detect a medium effect size (Cohen’s d = .50), and .92 power to detect a large effect size (d = .80) for alpha = 5%. Similarly, at the individual level with 20 trials, we had .56 power to detect a medium effect size (d = .50), and .92 power to detect a large effect size (d = .80).

## 3 Results

We examined children’s hemispheric activation upon performing two mental tasks, word generation or visuospatial memory. For each task after every trial, participants had to press a button to indicate whether they could generate many words, and whether the visual display was modified, respectively.

### 3.1 Behavioural responses

Behavioural performance on the visuospatial memory task was high (mean accuracy = 88%, range = [70%, 100%]). The percentage of trials for which they reported having thought of many words was somewhat lower (M = 77%, range = [45%, 100%]), and may have varied with the child’s understanding of “many” and/or their tendency to report their own performance as good or bad. As this was not a robust measure of behaviour (it was included only to encourage children to stay on task) this score was not considered further.

### 3.2 LI for each task: group level

Group level results are shown in Figure 3. We first illustrate the time course of the left and right hemispheres blood flow velocity for each task (Figure 3A and 3B). We then subtracted the right from the left activation, for each task, within their respective POI (Figure 3C and 3D). We calculated the significance of the LI for both tasks by comparing the left-right activation to 0. The LI for the word generation task was positive (M = 1.78 cm/s, SD = 3.33, 95% confidence interval (CI) = [.66, 2.90], Figure 3A and 3C) and significantly different from 0 (t_(19)_ = 3.33, p = 0.0035, Cohen’s d = .744) indicating left lateralisation at the group level. The LI for the visuospatial memory task was negative (M = −0.62 cm/s, SD = 1.57, 95% CI = [−1.36, 0.11]], Figure 3B and32D) but was not significantly different from zero in the time window of interest (t_(19)_ = −1.78, p = 0.0908, Cohen’s d = −.3983).

**Figure 3:**
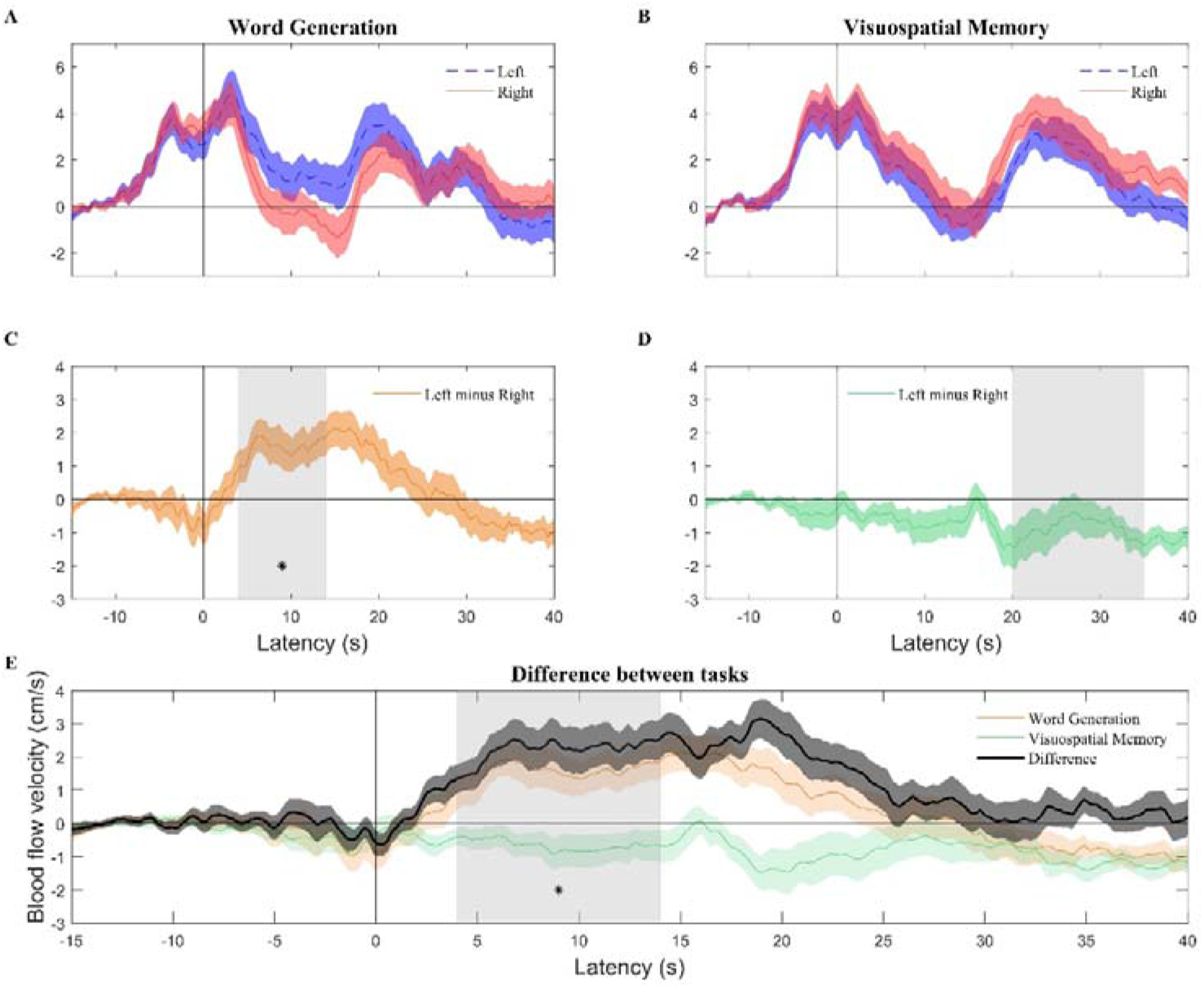
Grand average blood flow velocity for the left (dotted blue line) and right (solid red line) channels (**A**, **B**), and the left-minus-right difference (**C**, **D**) over time for the word generation (**A**, **C**) and visuospatial memory (**B**, **D**) task. **E** shows the left-minus-right difference (i.e., same as middle panels) for the word generation task (orange line) and the visuospatial memory task (green line), and the difference of these differences (black line). Grey areas indicate the periods of interest. Black asterisks indicate significant effects (p < .05).

### 3.3 LI for each task: individual level

We then examined the significance of LIs in individuals by comparing left-right differences to 0 in the language POI (4 to 14 s) for the language task and the visuospatial POI (20 to 35 s) for the visuospatial memory task. At the individual level (Figure 5), language was significantly lateralised to the left hemisphere for 50% of children (10/20), and to the right hemisphere for 5% of children (1/20). Visuospatial memory was significantly lateralised to the right hemisphere for 20% of children (4/20), and to the left hemisphere for 10% of children (2/20). The remaining participants did not show evidence of significant lateralisation. In addition, we examined the association between lateralisation for language and visuospatial memory. Although 9 of the 20 participants fell into the quadrant where they were numerically left lateralised for language and right lateralised for visuospatial memory, we found a significant correlation between the two functions (ρ = .48, p = 0.034). This indicated that participants with stronger left lateralisation for language also tend to have more leftwards lateralisation for visuospatial memory, and vice versa.

In addition, to check whether our approach to LI analysis made a difference to the lateralisation estimate from the empirical data, we ran a follow-up analysis using the statically biased peak selection method. Using this method, “significant” left lateralisation of language was still found in 50% of participants, but right lateralisation of visuospatial memory increased from 20% to 50% of participants. This confirms with that, indeed, peak selection can increase statistical bias towards reporting lateralisation.

### 3.4 LI difference between tasks: group level

At the group level, blood flow velocity for word generation was significantly different from the visuospatial memory task for the POI analysed (the language POI, Figure 3D). The word generation task was significantly more left-lateralized than the visuospatial memory task (word generation minus visuospatial memory = 2.21 cm/s, t_(19)_ =4.11, p<.001, Fig 3E).

### 3.5 LI difference between tasks: individual level

Finally, our main question was whether we could use our fTCD paradigm as an implicit measure of task-following in individual children. We assessed the sensitivity of detecting task-related hemispheric activation in individuals by comparing the left-minus-right differences between tasks. A significant effect of task was found in 55% (11/20) of our participants (see Figure 4), indicating clear evidence for task-following in just over half of individuals. In all the individuals with a significant difference between tasks, blood flow was more left-lateralised in the language task than in the visuospatial task.

**Figure 4.**
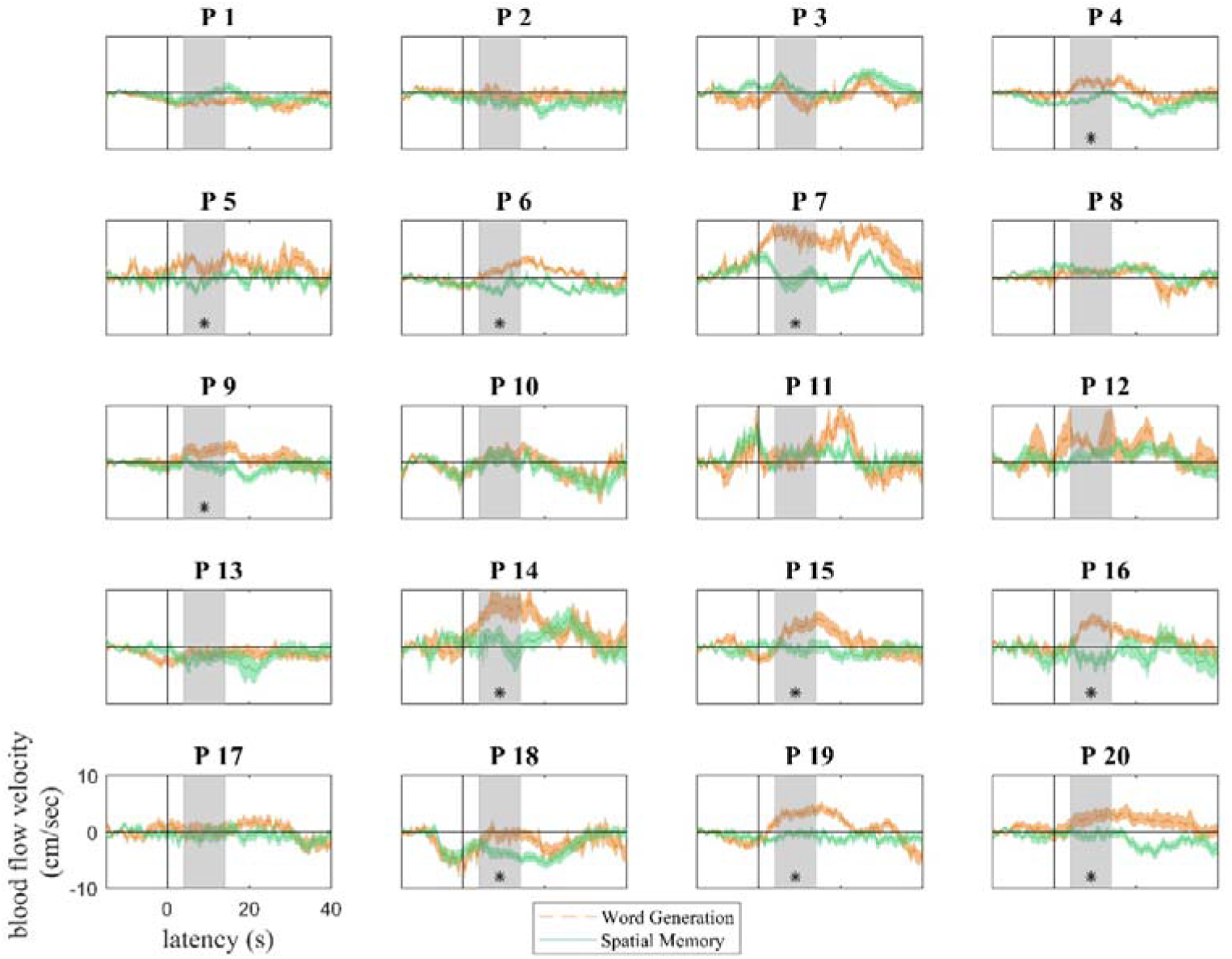
Individual participants pattern of activation (left-minus-right) for the word generation (dotted-orange line) and visuospatial memory (solid-green line) task, plotted ± standard error of the mean. Grey area indicates the period of interest for our analyses. Black asterisks indicate a significant difference in the POI. Eleven participants showed a statistically significant difference between the two tasks.

**Figure 5.**
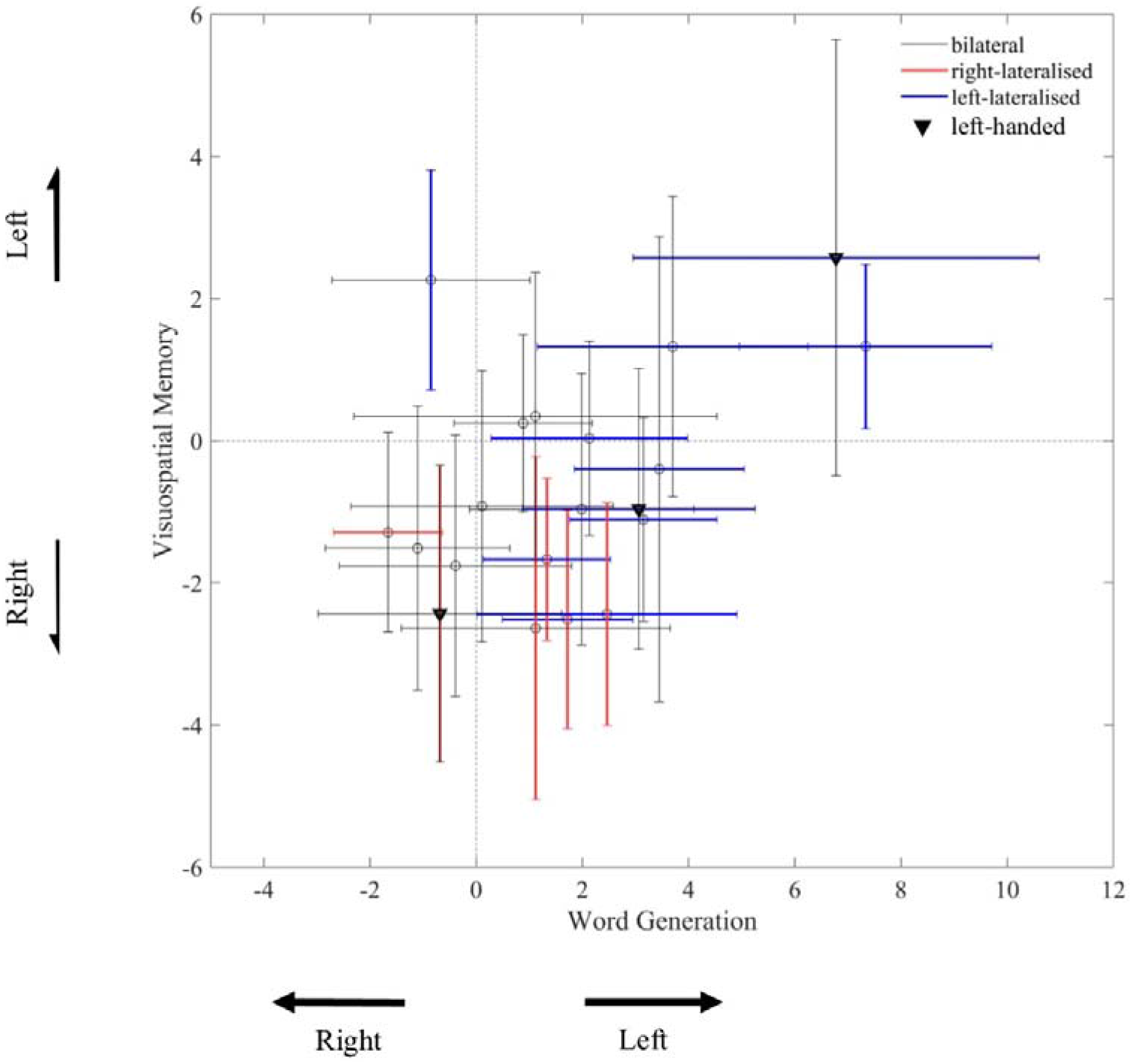
Scatterplot of laterality indices (LIs) of each participant for the word generation (POI = 4 to 14 s) and the visuospatial memory tasks (POI = 20 to 35 s), with 95% confidence interval for each participant (across trials). Participants with confidence intervals (CIs) overlapping zero are not considered to be lateralised (grey errors bars). Participants with CIs strictly < 0 are right-lateralised (red error bars), and participants with CIs strictly > 0 are left-lateralised (blue error bars). Left-handed participants are shown as black triangles.

## 4 Discussion

In this study we proposed a rigorous method for evaluating task-following in children based on the lateralisation of brain functions using functional transcranial Doppler ultrasound (fTCD). We designed a controlled, child-friendly paradigm in which children either silently generated words beginning with a particular letter (language task) or remembered the spatial location of letters on a screen (visuospatial task). We computed the left and right hemispheric blood flow velocity while children performed the tasks, and we inferred task-following from the difference in velocity between the two tasks. At the group level, we found significant evidence of task-following from the hemispheric activation, as seen by a significantly more leftward activation for the language task compared to the visuo-spatial memory task. This pattern was also found in 55% of individuals, indicating task-following in a subset of children. The null result in the remaining 45% of individuals is not directly interpretable and may reflect an absence of task-following, or a lack of sensitivity of our paradigm. In addition to these findings, we replicated previous literature in finding a significant left-lateralisation for the language task in children (Bishop et al., 2009; Groen et al., 2011, 2012). However, we did not observe the expected right-lateralisation for the visuospatial memory task, and we found less marked lateralisation of language and visuospatial memory in individuals, compared to what we expected from the literature.

The main aim of this study was to design a paradigm that could be used to assess task-following abilities in non-verbal individuals. To this end, we computed the patterns of left-minus-right hemispheric activation in response to a language task and a visuo-spatial memory task. A consistent difference in the hemispheric blood flow velocity between tasks, irrespective of the direction of this difference, would indicate the involvement of different brain activity in response to the two instructions, and thus indicate preserved task-following. However, even though we found a robust statistical difference in the activation for the tasks at the group level, we could only observe a statistical difference in 55% of individual participants. The current study used 20 trials per condition, in line with previous fTCD research (Badcock, Nye, et al., 2012; Groen et al., 2011; Whitehouse & Bishop, 2009). However, with 20 trials, we had only .56 power to detect a medium effect size (d=.50), in the individual subject analysis, so we may have failed to detect differences in the remaining individual children due to insufficient numbers of trials. More trials could be added by repeating some letters, and/or using the left-out letters that typically don’t allow generation of many words (e.g., X), but would necessarily increase the length of the experimental session, which may be challenging for some children.

A limitation of our approach was that we did not directly assess whether children were generating words in the language condition (e.g., by asking them to report the words generated). Our choice reflects our goal to apply this paradigm to non-speaking populations, for whom we cannot readily verify compliance through speech or behaviour. However, it does leave open the possibility that some children were not sufficiently engaged in the task or did not perform the task, despite self-reporting that they did so. Additionally, even though children reported the task to be engaging, it involved a long baseline period during which they were asked to clear their mind and "think of nothing". This might not be trivial, particularly for children, and it is possible that some participants engaged in language-related processes during the baseline period. In addition since we presented the same linguistic stimuli (an array of letters) across the two conditions, it is possible that children performed some linguistic processing when they were not instructed to or vice versa. This potentially reduces the difference in the activation between the tasks, making our test relatively conservative. Nonetheless it was essential to have identical stimuli to remove the visual confound that could otherwise drive different responses even in children who did not understand or perform the task. We instead mitigated the concern that children would perform the wrong task by using a blocked design, varying the condition every 10 trials, and observed a high degree of accuracy on the visual-spatial task. In the future, researchers may consider asking neurotypical children to report the words they generated, to examine the contribution of task compliance to individual differences in lateralisation. However, a previous study on language generation in adults found no difference in lateralisation between overt and covert word generation (Gutierrez-Sigut et al., 2015), implying that overt word generation is not necessary for left-hemisphere activation. Despite these limitations, which prevent us from drawing any conclusions in children that do not show differential patterns of lateralisation, the positive results found in half of the participants is objective evidence for intact task-following in these children. For future clinical use, the limitations presented here need to be addressed in order to establish a paradigm that shows a higher sensitivity to detect effects in neurotypical children.

In addition to our main interest in task-following, we also reported the lateralisation of each task separately, using statistically robust analyses to calculate LIs. As expected from the literature, we found significant left-lateralisation of language at the group level. However, at the individual level, the lateralisation was not as pronounced as expected. Only 50% (10/20) of children had significant left-lateralisation of language, and 25% (5/20) had significant right-lateralisation of visuospatial memory. This rate is lower than previously reported, i.e., around 70% of people being left-lateralised for language (e.g., Knecht et al., 2000; Lust, Geuze, Groothuis, & Bouma, 2011; Whitehouse & Bishop, 2009), and 70% of people being right-lateralised for visuospatial memory (e.g., Groen et al., 2012; Whitehouse & Bishop, 2009). The numerically lower lateralisation found in our study may reflect the use of identical stimuli between tasks. It is possible that the identical stimuli tended to encourage children to perform the incorrect task (as above), and/or it is possible that part of the differential lateralisation observed in previous work is driven by the confounding differences in visual stimuli. Our data confirm that there is at least some lateralisation of these processes even when visual stimuli are closely controlled.

A separate explanation for our observation of relatively low lateralisation in individuals is that many previous analyses may have been statistically biased towards detecting lateralisation. Many fTCD studies derive LIs by defining a time-window around a peak in the left-minus-right blood flow velocity, and then analysing the data within this time-window (Deppe et al., 2004). As we showed with simulation (see Methods), this peak selection technique introduces a bias if these data are subsequently compared to zero at the individual or group level, to infer lateralisation (a technique known as double dipping). In particular, it increases type I error rate (probability of incorrectly rejecting the null hypothesis when it is true, i.e., false positives). It also artificially creates bimodality in the distribution of LIs in the population (Woodhead et al., 2018). In this study, we overcame this problem by computing the difference in the left-right blood flow velocity difference over an entire pre-defined POI (as introduced by Woodhead et al., 2018), which, as our simulations showed, brings the type I error back to the scientific standard of 5%. The peak selection method may still be suitable to compare lateralisation between tasks or between groups, but should not be used to test whether individual or group lateralisation is different from chance, because the multiple comparisons inherent in selecting the peak from continuous data have not been accounted for. Had we used the peak selection method in our study, we would have increased our rate of “significant” right-lateralisation for the visuo-spatial task from 20% to 50% of participants. However, this does not entirely account for the difference that we observe compared to previous literature, suggesting that at least some of the difference between our results and previous literature may be due to differences in the stimuli used or population examined. It is possible that the lateralisation of children age 9-12 is not as strong as previously suggested, although significant lateralisation was still present in some individuals even with statistically robust methods. Our sample size was small (n=20) and a larger study with appropriate statistics would be needed to give a reliable estimate of the population’s lateralisation.

Upon analysing the association between the lateralisation for the two tasks in individuals, we found a significant positive correlation between language and visuospatial memory lateralisation. In other words, despite the group-level typical left lateralisation for language and right lateralisation for visuospatial memory, individuals who were more left lateralised for language were also more leftwards lateralised for visuospatial memory. This correlation is consistent with previous reports (Flöel et al., 2005; Whitehouse & Bishop, 2009). It can be taken as evidence against a causal view of hemispheric specialisation in which localisation of one function to one hemisphere causes localisation of the other function to the other hemisphere (e.g., language is left lateralised *because* visuospatial memory is right lateralised; Whitehouse & Bishop, 2009; Cai et al., 2013). However, the data are also not well explained by the dominant alternate view, in which hemispheric lateralisation of each function is independent (Bryden et al., 1983), as this predicts no association in LI between tasks. Instead, our data suggest an *association* in which individuals who tend to rely more on their left hemisphere in one task, will also tend to rely more on this hemisphere in the other task. Further work is needed to understand the extent to which this reflects the tendency for participants to engage in some language processing in the visuo-spatial memory task and vice versa, particularly since it has also been observed in paradigms that use different materials between tasks (e.g., Flöel et al., 2005; Whitehouse & Bishop, 2009), and seems at odds with the prevailing view that lateralisation of these two tasks should be complimentary.

## 5 Conclusion

We measured brain activation in children using a portable and inexpensive neuroimaging device, fTCD. We analysed lateralisation of neural blood flow in response to a language task and a visuospatial memory task performed on identical visual stimuli. Two main findings emerge. First, we were able to observe task-following from the brain data of just over half the participants, making our results a promising basis for future clinical tests. By analysing the hemispheric activation pattern across the two tasks, statistically robust differences were observed in 55% of individual children. Second, our results indicate that the lateralisation of neurotypical children may not be as pronounced as previous research suggests. While previous fTCD research found left-lateralisation of language in about 70% of children, we were only able to see this pattern in 50% of our participants. Similarly, typical fTCD research found right-lateralisation of visuo-spatial memory in 70%, while we found this pattern in 20% of our participants. This was potentially due to controlled stimulus presentation, a tendency for children to perform some of the opposite task even when not instructed to, and/or less biased statistical assessment of lateralisation. Overall, our methods constitute a promising step towards the neural measurement of task-following abilities in children. These methods, however, need further development before they can be used as an assessment tool in special populations, possibly with more trials, an independent index of subject compliance, and refined paradigms to maximally differentiate between the two hemispheres.

## Declarations of interest

none

## Funding

This work was supported by a Cognitive science Postgraduate research Grant from the Department of Cognitive Science at Macquarie University. AW was funded by an ARC Future Fellowship (FT170100105) and MRC (U.K) intramural funding SUAG/052/G101400.

## Acknowledgements

We thank Martyn Churcher and Kristie Tainton for their recordings of the pirate voices.

## Footnotes

1 We use ‘identify-first’ language (‘autistic person’) rather than person-first language (‘person with autism’), because it is the preferred term of autistic activists (e.g. Sinclair, 2013) and many autistic people and their families (Kenny et al., 2016) and is less associated with stigma (Gernsbacher, 2017).

## Notes

### Competing Interest Statement

The authors have declared no competing interest.

